# Multiplexed Short-wave Infrared Imaging Highlights Anatomical Structures in Mice

**DOI:** 10.1101/2024.01.29.577849

**Authors:** Xingjian Zhong, Amish Patel, Yidan Sun, Alexander M. Saeboe, Allison M. Dennis

## Abstract

While multiplexed fluorescence imaging is frequently used for *in vitro* microscopy, extending the technique to whole animal imaging *in vivo* has remained challenging due to the attenuation and scattering of visible and traditional near infrared (NIR-I) wavelengths. Fluorescence imaging using short-wave infrared (SWIR, 1000 – 1700 nm, a.k.a. NIR-II) light enables deeper tissue penetration for preclinical imaging compared to previous methods due to reduced tissue scattering and minimal background autofluorescence in this optical window. Combining NIR-I excitation wavelengths with multiple distinct SWIR emission peaks presents a tremendous opportunity to distinguish multiple fluorophores with high precision for non-invasive, multiplexed anatomical imaging in small animal models. SWIR-emitting semiconductor quantum dots (QDs) with tunable emission peaks and optical stability have emerged as powerful contrast agents, but SWIR imaging demonstrations have yet to move beyond two-color imaging schemes. In this study, we engineered a set of three high quantum yield lead sulfide/cadmium sulfide (PbS/CdS) core/shell QDs with distinct SWIR emissions ranging from 1100 – 1550 nm and utilize these for simultaneous three-color imaging in mice. We first use QDs to non-invasively track lymphatic drainage, highlighting the detailed network of lymphatic vessels with high-resolution with a widefield imaging over a 2 hr period. We then perform multiplexed imaging with all three QDs to distinctly visualize the lymphatic system and spatially overlapping vasculature network. This work establishes optimized SWIR QDs for next-generation multiplexed preclinical imaging, moving beyond the capability of previous dual-labeling techniques. The capacity to discriminate several fluorescent labels through non-invasive NIR-I excitation and SWIR detection unlocks numerous opportunities for studies of disease progression, drug biodistribution, and cell trafficking dynamics in living organisms.

## Introduction

Fluorescence imaging is a widely adapted bioimaging technique owing to its non-ionizing radiation and relative low cost, but deep tissue *in vivo* imaging is challenging due to the opaque nature of skin and tissue. Researchers circumvent these issues by imaging in near-infrared (NIR) optical tissue windows with NIR-emitting contrast agents^1,2^ or multi-photon excitation in the NIR to excite bluer emitters.^3,4^ In the NIR-I window (650 - 900 nm), hemoglobin absorbance and tissue scattering are lower than in the visible wavelength regime (490-650 nm).^5^ However, tissue autofluorescence and tissue scattering still limit applications such as anatomical imaging or quantitative targeted imaging.^6,7^ To further advance fluorescence imaging, NIR-II imaging, also named short-wave infrared (SWIR) fluorescence imaging, has emerged as an *in vivo* bioimaging technique that utilizes the biological windows from 1000 – 1700 nm.^8,9^ Compared to conventional NIR-I window, the SWIR biological window exhibits reduced tissue scattering and low autofluorescence to enable deeper tissue penetration with high signal-to-noise imaging contrast; light absorption by the tissue is relatively low, except for a large water absorption peak ∼1400 nm.^10,11^ The ongoing development of SWIR-emitting contrast agents is critical to the continued advancement of fluorescence imaging in this longer wavelength range. Following the initial demonstration of biomedical imaging in the SWIR using carbon nanotubes^7^, *in vivo* SWIR imaging has been demonstrated with fluorescent contrast agents ranging from semiconductor quantum dots (QDs)^12–15^, small molecule dyes^16– 20^, lanthanide-doped nanoparticles^21–23^, and engineered fluorescent proteins.^24^

The improved image quality enabled by SWIR imaging allows for *in vivo* deep tissue multiplexing, which has been commonly used in cellular and tissue imaging to highlight different targets of interest in the same specimen.^25^ With low background, high contrast, and high resolution compared to widefield NIR-I imaging, multiple fluorophores can be distinguished with high precision, opening tremendous opportunities to study disease models, new biological phenomena, and quantitative *in vivo* sensing.^26^ Different approaches are needed for multiplexed SWIR imaging depending on the contrast agents being used. Multiplexing with SWIR emitting dyes is achieved using discrete excitation wavelengths to distinguish the fluorescence from different emitters, which is all collected using the same long-pass emission filter.^27–29^ The emission peaks of the fluorescent dyes and a newly developed fluorescent protein, however, are typically in the NIR-I; SWIR imaging of these emitters relies on collecting photons from the long wavelength tail of the emission peak, precluding spectral separation of the emission peaks and relying on the dimmest portion of the contrast agent emission. Another study engineered lanthanide-doped nanoparticles with distinct luminescence lifetimes for signal separation via time-gated imaging.^30^ In comparison, there are a handful of examples where QDs have been paired with another QD or dye for multiplexed imaging using a single excitation wavelength and camera filter sets for spectral separation.^31–33^

With few reports of multiplexed SWIR imaging, and those almost exclusively exploring dual-color imaging, we explore three-color *in vivo* SWIR imaging in live mice. In this study, we expanded the synthesis of high quality PbS/CdS core/shell QDs and selected three QDs with distinct emission profiles in the SWIR imaging window. Using an IR VIVO preclinical live animal imager with grating-based hyperspectral and filter-based multispectral imaging capabilities, we non-invasively visualize the lymphatic pathway in high resolution and monitor the lymphatic drainage over time. Then using three colors, we map out the lymphatic system and spatially overlapped vascular structures simultaneously, demonstrating the potential of multiplexed SWIR imaging using semiconductor QDs.

## Results

### Synthesis of PbS/CdS QDs across SWIR window

The synthesis of PbS/CdS core/shell QDs proceeds in three steps: *1)* synthesis of PbS core QDs, *2)* cation exchange of Pb for Cd to form a CdS shell, and *3)* coating of the particles to render them water soluble (**Fig. 1A**). We synthesized three different PbS cores via hot-injection precipitation using lead chloride (PbCl_2_) and sulfur in oleylamine (OlAm) as the lead and sulfur precursors, respectively. Core sizes were controlled by modifying reaction time and temperature, whereby increasing either results in larger particles with a first excitonic (1s) absorption feature at longer wavelengths. Since the reaction occurs at relatively low temperature, quenching the reaction with a liquid nitrogen-chilled hexane/ethanol mixture to simultaneously stop the reaction and precipitate the particles was key to maintaining sample homogeneity. Additionally, for the small core synthesis, we also reduced precursor concentrations to increase the particle homogeneity at the smaller size. The absorbance of three synthesized cores shows distinct and narrow 1s features at 1220 nm, 1370 nm and 1575 nm for small, medium and large cores, respectively (**Fig. 1B**). TEM images and small angle X-ray scattering (SAXS) measurements show the size, morphology, and homogeneity of these PbS QDs with diameters of 5.71 nm, 6.29 nm, and 7.41 nm, respectively (**Fig. S1 and S2, Table S1**). Similar to previous reports,^34,35^ we found that these OlAm-capped QDs were not stable after cleaning through precipitation and therefore not suitable for long-term storage. We thus performed ligand exchange to replace OlAm with oleic acid (OA) to increase the stability of the cores.

**Figure 1.**
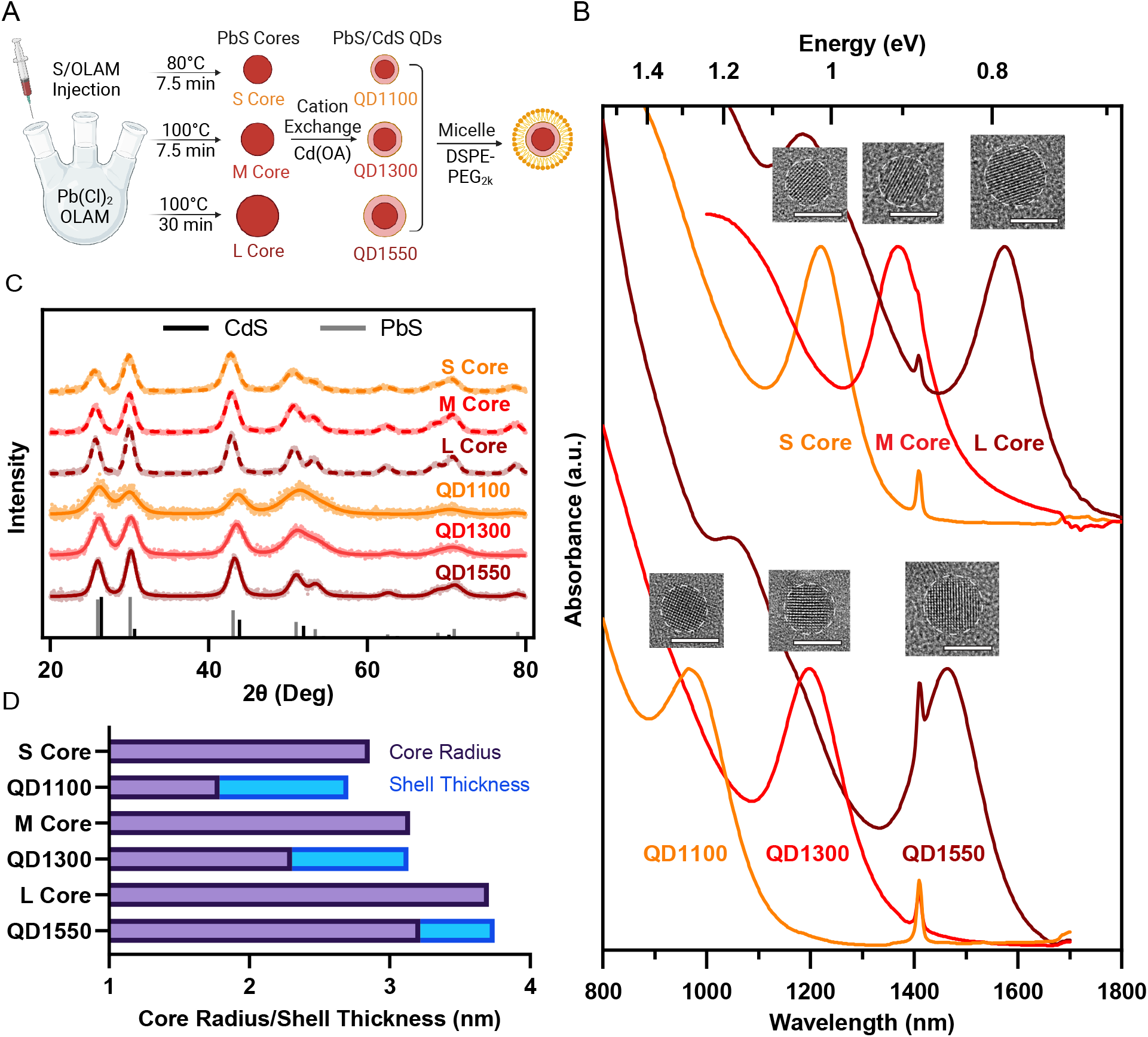
QD synthesis and characterization. A) Schematic of nanoparticle preparation including core synthesis, cation exchange-based shell formation, and micelle encapsulation. (Created with BioRender.com). B) Absorbance spectra of PbS cores and PbS/CdS core/shell QDs shown with representative TEM images (scale bar = 5 nm). Note: Peak at 1400 nm is an artifact of TCE (solvent) absorption. C) XRD profiles of PbS cores and PbS/CdS core/shell QDs with reference CdS peaks in black (ICDD: 01-089-0440) and PbS peaks in grey (ICDD: 01-077-0244). D) Core radius and shell thickness measurements for each QD ensemble, calculated by combining the size measurements from SAXS and elemental analysis.

We empirically determined cation exchange conditions with cadmium oleate to convert the three PbS cores to PbS/CdS core/shell QDs with 1s peaks at 965 nm, 1200 nm, and 1465 nm, named QD1100, QD1300, and QD1550, respectively, in reference to their photoluminescence emission peak wavelength positions (**Fig. 1B, Table S1**). In the cation exchange reaction, the size of the particles remains constant while cadmium atoms replace the surface lead atoms to form a CdS shell layer. This leads to a blue-shift of the 1s peaks through shelling despite similar sizing results to the cores in SAXS measurements (**Fig. 1B**,**C; Fig. S2**). Structural characterization with X-ray diffraction (XRD) shows only PbS peaks in the core samples, while the merged peak around 53° indicates the presence of both PbS and CdS in the core/shell samples (**Fig. 1C**). An increase in the particle crystal size is also indicated through XRD peak narrowing from the small to large core. Using microwave plasma atomic emission spectrometry (MP-AES) measurements for elemental analysis, we combined the ratio of Pb and Cd in each core/shell QD sample with sizing information to estimate the core radius and shell thickness in each of the particles. As expected, higher temperatures during the QD1100 and QD1300 cation exchange reactions yielded thicker shells than the lower temperature reaction used to generate QD1550. These QDs exhibit emission peaks at 1115 nm, 1310 nm, and 1545 nm with absolute quantum yields (QY) of 21.6%, 23.4% and 18.0+% (measured up to the 1700 nm detector limit), respectively (**Fig. 2A**).

**Figure 2.**
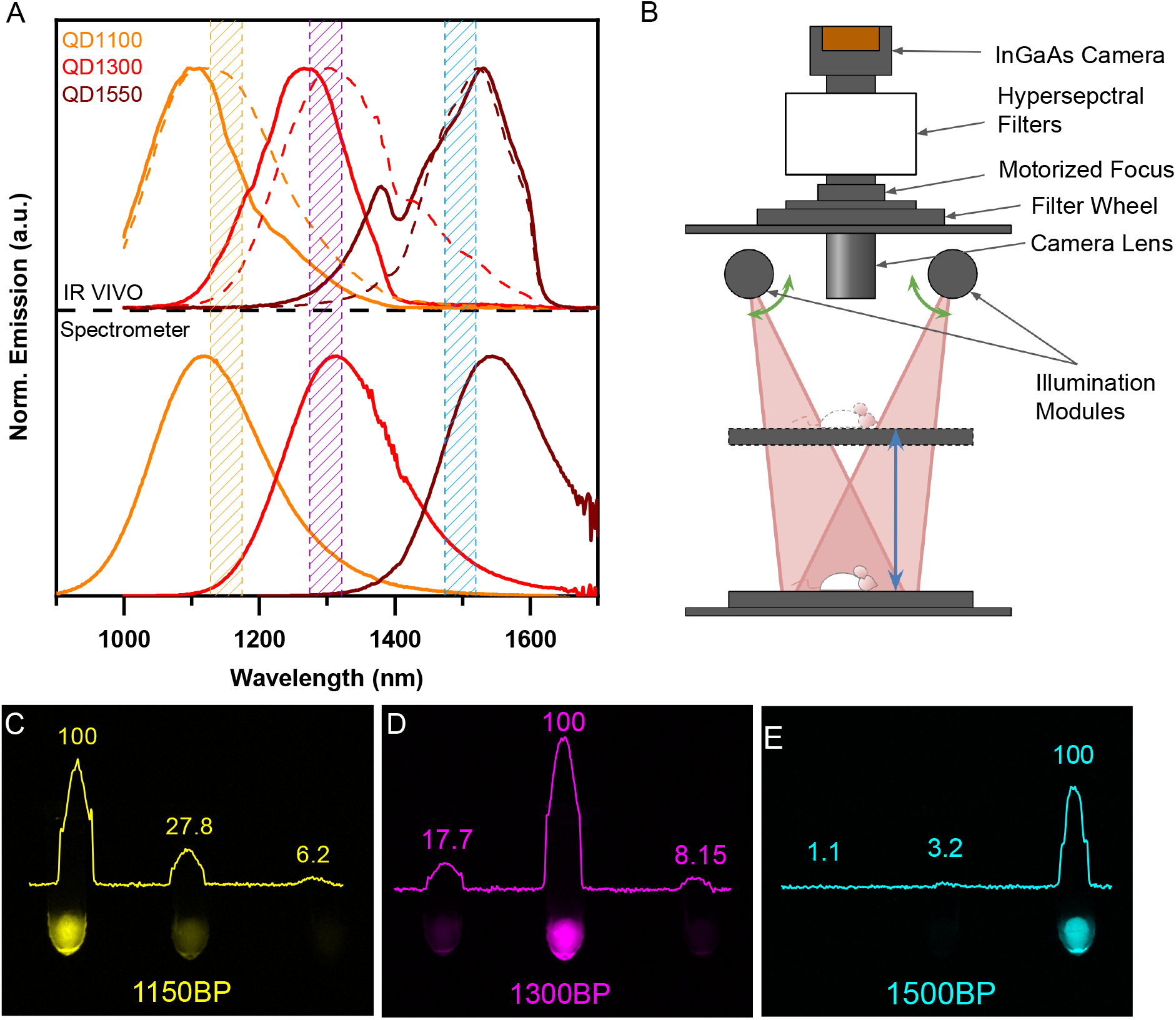
Optical properties of the QDs measured with SWIR imager and spectrometer. A) Emission profiles of QD1100, QD1300, and QD1550 PbS/CdS QDs in aqueous solution (solid line) and in solvent-free dropcast samples (dashed line) measured with the IR VIVO imager (top) and trichloroethylene (TCE)- suspended particles measured with a spectrometer (bottom). Spectral location of three bandpass filters used later in the study are shown as boxes filled with diagonal lines. B) Schematic of the key components of IR VIVO (Photon Etc.) SWIR imager includes a multi-laser illumination module, filter wheel for filter combinations, hyperspectral filter cube, and an InGaAs camera for signal acquisition. C – E) Three tubes containing (from left to right) QD1100, QD1300, and QD1550 imaged with 1150BP, 1300BP, and 1500BP filters demonstrate the extent of crosstalk for each emitter in the different bandpass (BP) filters. Images are pseudo-colored yellow, magenta, and cyan. Intensity linescans are normalized to the QD ensemble with maximum intensity, and the relative intensities of the other tubes is indicated in the images. Excitation at 808 nm; integration time: 0.02 s.

Following the successful synthesis of PbS/CdS core/shell QDs, we encapsulated the particles in biocompatible PEGylated phospholipid micelles for imaging. All PbS/CdS QDs were coated with 1,2-distearoyl-sn-glycero-3-phosphoethanolamine-N-[amino(polyethylene glycol)-2000] (DSPE-PEG_2k_), resulting in three QD samples with similar hydrodynamic diameter and surface charge (**Fig S3)**. Encapsulated particles were ultracentrifuged to remove empty micelles, and the concentration of the particles was measured with MP-AES. After aqueous transfer, we measured the photoluminescence of the particles in the IR VIVO imager in hyperspectral mode to compare with the particle photoluminescence emission spectra in organic solution prior to the encapsulation (**Fig. 2**). The loss of signal at 1600 nm in the IR VIVO is attributed to the detector cutoff. By comparing solvent-free dropcast samples measured with the IR VIVO, which exhibit photoluminescence spectra similar to the emission profiles of the trichloroethylene (TCE)-suspended samples measured in a benchtop spectrometer, to the micelle-encapsulated QDs imaged in hyperspectral mode in aqueous solution, we can see the impact of water absorbance on the photoluminescence spectra. Specifically, the onset of water absorbance around 1125 nm decreases the emission intensity of QD1100 above 1125 nm. The increase in water absorption above 1300 nm effectively blue-shifts the QD1300 peak after normalization and quenches the long-wavelength emission tail. The intensity of water absorption from 1400-1500 nm is seen as a dip in the QD1500 emission peak as well as a general attenuation of the brightness from this emitter. Peak brightness-matched QD solutions were imaged with six BP filters to test their crosstalk in each channel (**Fig 2C – E, Fig. S5**). We noticed unexpectedly high crosstalk in the 1100BP channel from QD1550, which is due to the filter ineffectively blocking light >1500 nm. Comparing the crosstalk between the rest of the channels, we decided to mainly utilize 1150BP, 1300BP, and 1500BP filters for the subsequent multiplexed imaging.

### Imaging lymphatic drainage over time

We first visualized lymphatic drainage in a live mouse using QD1550. We used an injection site at the mouse tail base that would reportedly highlight drainage from the injection site to the inguinal-axillary pathway;^36–38^ however, none of the previous studies had the imaging capacity to map the full drainage network in a live mouse. In this experiment, 50 μL of 8μM QD1550 in sterile saline were subcutaneously injected midline at the base of the tail, and imaging began about 3 min post injection (p.i.). We imaged the mouse with 808 nm laser excitation every 90 s from 3 min to 35 min p.i. and from 90 min to 126 min p.i. using a 1250 nm long-pass (1250LP) filter and the 1550BP filter. We successfully produced a clear map of lymphatic drainage including a tortuous network of thin lymphatic vessels from the injection site primarily to the left inguinal lymph node (LN) and both left and right axillary LNs (**Fig. 3A – B**). Comparing the two filters, we see that the 1550BP filter images exhibit better imaging resolution, improving visualization of fine lymphatic vessel structure (**Fig. 3C**). When analyzed quantitively across region-of-interest (ROI) lines in the images, we could identify thinner vessels with finer resolution with the 1550BP filter; for example, one vessel appeared to be 376.4 μm wide when imaged with the 1550BP filter, while the same vessel imaged with the 1250LP filter was 593.7 μm wide (**Fig. 3D – E**). Systematically comparing all the feature widths along the ROI demonstrates that the FWHM of peaks measured with the 1550BP filter were consistently thinner with a cluster of features ∼ 380 μm wide in an imaging system with a camera pixel size of 125 μm (**Fig. 3F**). Since the signal to noise ratio was comparable when imaging with the 1550BP or 1250LP (**Fig. 3D**), we suspect the differences were caused by chromatic aberrations that are more apparent when imaging across the wide range of wavelengths covered by the LP filter in comparison to the short bandpass range as well as the 1550BP imaging benefiting from water absorption attenuating scattered light within the sample, which was also observed by another study to increase imaging contrast at 1450 nm.^39^ As the narrower BP filter only collects a portion of the light emitted by the QDs, imaging with the BP filter requires a longer exposure time to collect comparable signal to the LP filter. Specifically, we used 5× longer exposure time with the BP filter (0.5 s vs. 0.1 s) to collect roughly equivalent raw signal counts with the LP filter.

**Figure 3.**
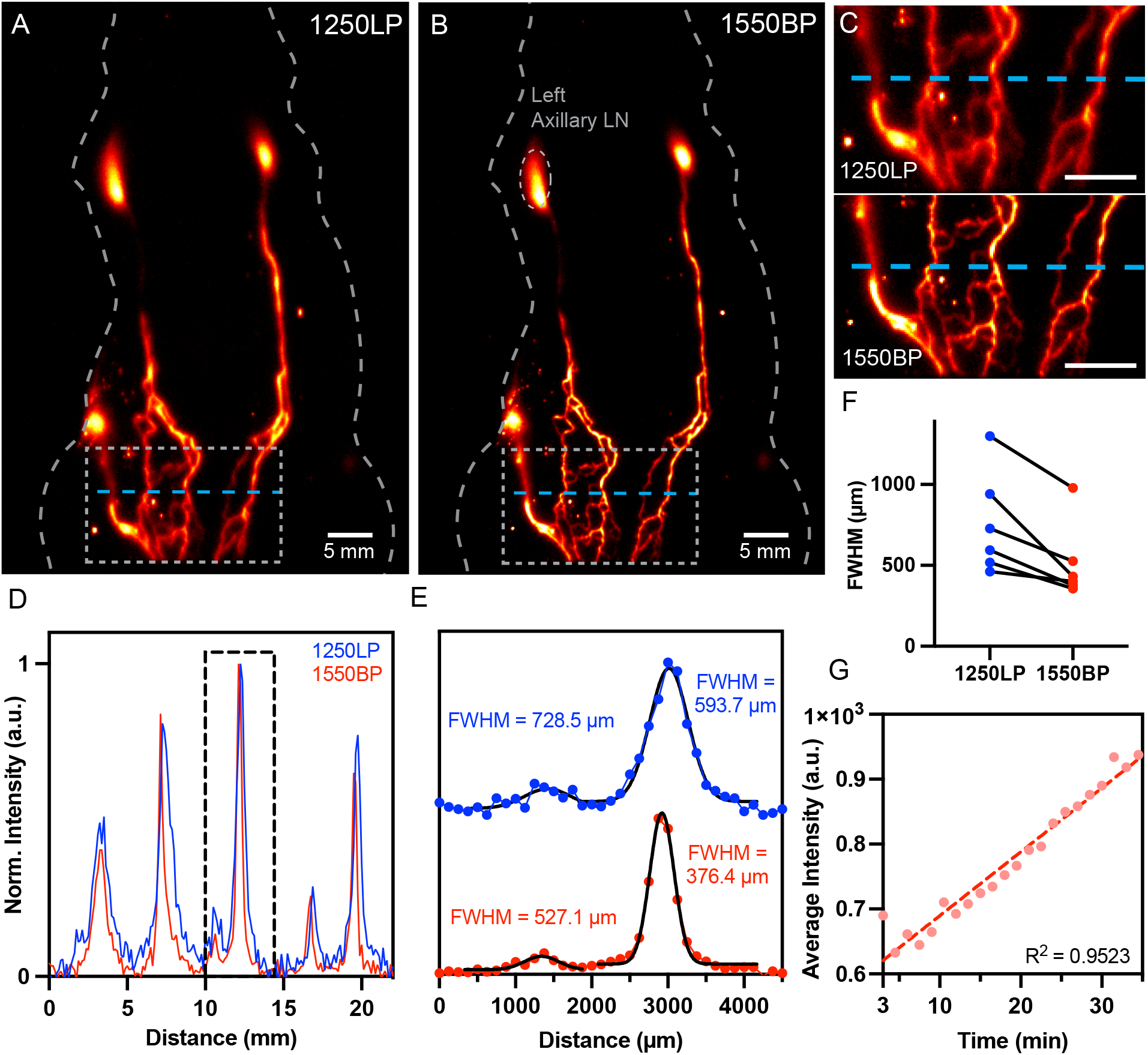
Single color imaging of lymphatic drainage from tail base subcutaneous QD injection. A) 1250LP and B) 1550BP images 100 minutes post subcutaneous injection. C) Zoom-in comparison of the rectangular ROIs in A and B (scale bar = 5 mm). D) Average signal intensity changes with imaging time from the left upper axillary lymph node (drawn in B) calculated from 1550 BP images. E) Normalized signal intensities across the ROI lines (blue dashed) drawn from the same location on mouse in C. F) Zoom-in comparison of the signal intensities in the black ROI region from E. G) Comparison of all peak FWHMs from E extracted using Gaussian fits.

By analyzing time lapse images, we semi-quantitatively track the drainage and accumulation in LNs using the average or summed signal counts at specific ROIs. For example, when plotting the average signal in the left axillary LN over time, we observe a linear increase in the intensity during the first imaging session, which lasted from 3-35 min p.i. (**Fig. 3G)**. This indicated a steady drainage rate over this period. During the second imaging session, which encompassed 90–126 min p.i., we observe a decreasing rate of accumulation in the axillary LN followed by a plateau in the photoluminescence intensity (**Fig. S5**). This indicates that most of the particles moving through the lymphatic network drained to corresponding LNs within 2 hr of the injection, and the drainage rate has decreased significantly. It should be noted that a substantial change in mouse position precludes quantitative comparisons between imaging sessions due to changes in the depth of the anatomical structure impacting raw signal intensities. Therefore, we separate the analysis of the two imaging sessions, focusing on comparisons drawn while the mouse remained in a relatively still position. Using SWIR imaging, we demonstrated the detailed mapping of lymphatic structures and provided a semi-quantitative method to monitor the lymphatic drainage of contrast agents.

### Multiplexed three color imaging to highlight anatomical structures

With the lymphatic pathway identified, we multiplexed three QDs to image overlapping lymphatic and vascular structures. As illustrated in **Figure 4A**, QD1100 and QD1300 were first subcutaneously injected into the left and right sides of the tail base, respectively, to highlight two sides of the lymphatic network. QD1100 are clearly visible on the right side of the picture, draining from the injection site to the axillary LN, while QD1300 drained through the inguinal LN to the axillary LN (left side of the picture). Prior to injecting QD1550, we observed no crosstalk from QD1550 in the 1100 nm BP filter channel, and no signal in the 1500BP and 1550BP filter channels (**Fig. S6**). QD1550 were retro-orbitally (r.o.) injected 2 hr after the initial subcutaneous injections and additional images collected (**Fig. S7**). In the 1100BP filter channel, the unfocused bleed-through signal from QD1550 was obvious, as was also seen with the images of the QD solutions (**Fig. S4**). Vascular circulation was seen in the 1500BP and 1550BP channels, highlighting detailed vessel structures with some liver accumulation. We selected the 1150BP, 1300BP, and 1500BP filters for visualizing the multiplexed imaging of the three QDs (**Fig. 4**). Signal from QD1550 was brighter than the other two QDs due to enhanced delivery efficiency through i.v. injection compared to partial drainage of contrast agent in the lymphatic system. This enhanced brightness exacerbated crosstalk especially in the liver region in the 1300BP channel. However, the fine anatomical structures of the lymphatic network and the vascular structure are still easily identifiable in the 3-color composite image. When examining the two linear ROIs drawn in the composite image, the two axial lymph nodes with distinct signals from QD1100 and QD1300 are clearly visible with intensity spikes from vascular structures clear in the 1550BP channel, including a narrow vascular structure overlapped with the right axillary LN around 2.9 mm. In ROI-2, each detailed vessel appears as a small intensity peak in the 1500BP from QD1550, and the lymphatic vessel and blood vessel near position 2800 μm can be distinguished despite their close proximity. The inguinal LN pictured on the left is so brightly illuminated by QD1300 that it saturates the signal in the 1300BP channel around 400 μm.

**Figure 4.**
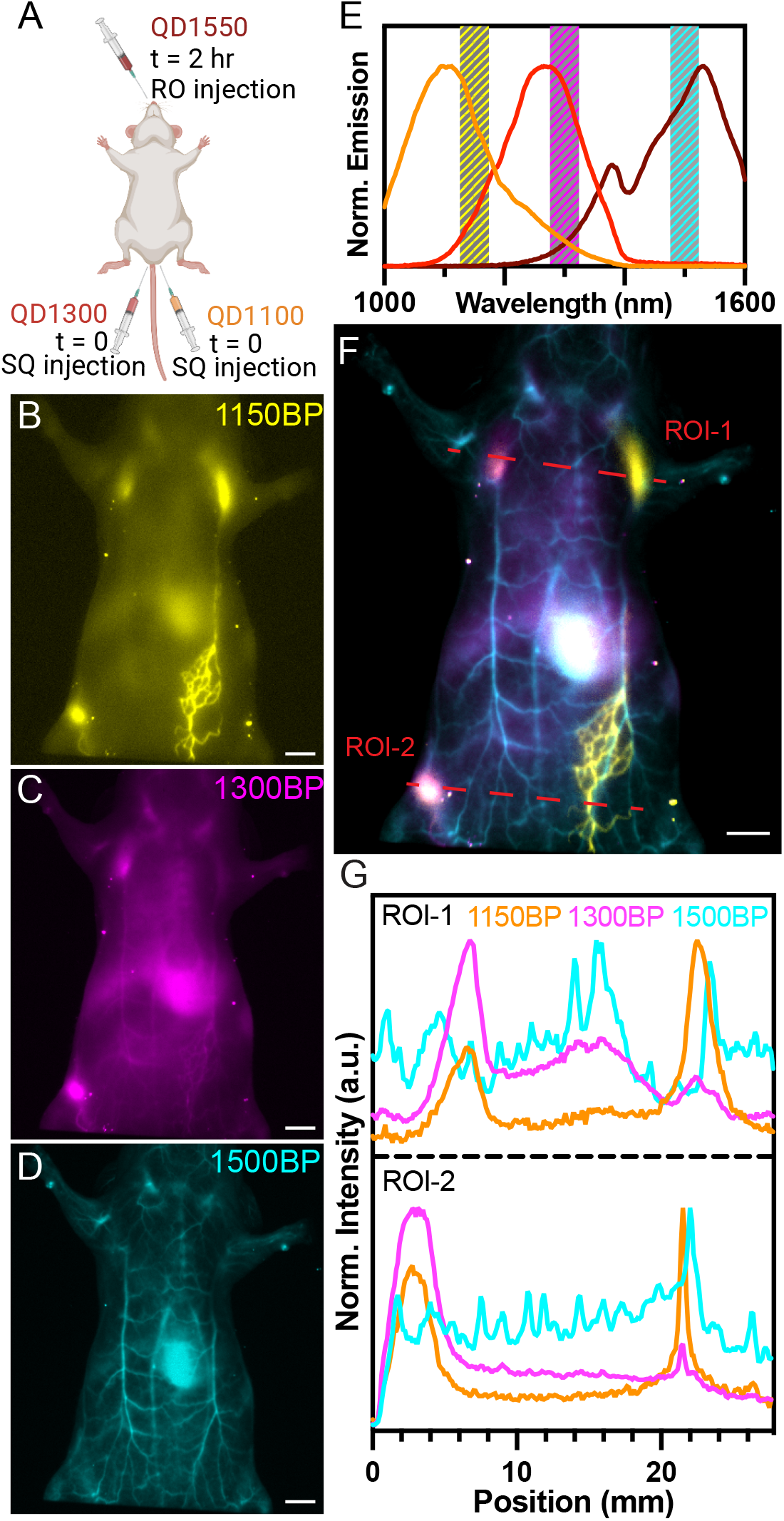
Three-color multiplexed mouse imaging. A) Experimental scheme including injection location and timing information. (Created with BioRender.com) B – D) Mouse imaged after subcutaneous injection of QD1100 and QD1300 on the left and right sides of the base of the tail, respectively, and retroorbital injection of QD1550 with 1150BP, 1300BP and 1500BP filters (scale bar = 5 mm). Images are pseudo-colored yellow, magenta, and cyan, respectively. E) Emission spectra of QDs and the three corresponding bandpass filter locations. F) Composite of images B – D in their corresponding colors (scale bar = 5 mm). G) Normalized signal intensity with respect to position along the two ROI lines drawn in F.

## Discussion

Multiplexed fluorescence imaging, which leverages spectral resolution to distinguish multiple fluorophores in a single specimen, provides an attractive platform for continuous *in vivo* tracking of biological structures and processes.^25^ However, previous attempts at multiplexed animal imaging were often confounded by background autofluorescence and limited tissue penetration depth in the visible and conventional near infrared (NIR-I) spectral range. These limitations hinder quantification and often require complex algorithms to differentiate closely spaced emission signals.^40,41^ The recent emergence of SWIR fluorescence imaging has helped overcome such issues by capitalizing on an optical window from 1000-1700 nm, where scattering and autofluorescence are minimized, and narrower windows enjoy greatly reduced absorbance from tissue components. Utilizing these optical tissue windows enables centimeter deep imaging and micrometer-range spatial resolution.^42^ This enables multiplexing with much higher signal-to-noise for straightforward fluorophore differentiation. Nonetheless, current examples of SWIR multiplexing remain limited to dual-color imaging, ^28,29,31,33^ and none uses multiple fluorophores with discrete SWIR emission peaks to differentiate signals based solely on emission wavelength.^27,30^

To push the boundaries of SWIR multiplexing, appropriate contrast agents must exhibit narrow emission bandwidths for spectral separation and relatively high and balanced brightness across labels to simplify dosing and signal discrimination. Semiconductor QDs stand out as an option due to their high quantum yield, stability, and tunable SWIR emission, making them ideal candidates for multiplexing.^11–13^ Moreover, distinct colors of QDs can be interchangeably incorporated into coatings and/or drug delivery vehicles, minimizing variation in imaging outcomes related to the physiochemical properties of different contrast agents such as size, hydrophobicity, charge, and surface properties. Here, we used PbS/CdS core/shell QDs due to their wide range of tunable emissions across the SWIR imaging window. CdS shell enhances brightness by reducing surface defects and serving as a protective layer against oxidation. While PbS/CdS QDs have been demonstrated as contrast agents for SWIR imaging, no study has optimized them for multicolor multiplexed imaging. After synthesis, all QDs were encapsulated with the same PEGylated phospholipids to ensure uniform size and surface coating. The final constructs ranged from 10 – 100 nm to enable efficient lymphatic drainage.^43^

In summary, we successfully engineered three PbS/CdS core/shell QDs with distinct SWIR emission profiles ideal for multiplexed fluorescence imaging. We successfully demonstrated the non-invasive observation of lymphatic drainage of semiconductor QDs revealing a tortuous network of lymphatic vessels using live, whole-body fluorescence imaging in mice. Time lapse imaging over 2 hr depicted the accumulation of nanoparticles in the axillary LN and changes in the drainage rate over time. We then demonstrated simultaneous three-color mapping of the lymphatic system and vasculature. SWIR imaging provided excellent contrast to identify structures and quantify features down to ∼350 μm without additional signal processing. This work establishes micelle-encapsulated SWIR QDs for straightforward multiplexing and non-invasive *in vivo* anatomical imaging.

## Methods

### Materials

Lead chloride (PbCl_2_, 98%), oleylamine (OlAm, technical grade, 70%), sulfur (S, trace metal grade), oleic acid (OA, technical grade, 90%), cadmium oxide (CdO, trace metal grade), 1-octadecene (ODE, technical grade, 90%), trichloroethylene (TCE, 99.5%), and hexane (anhydrous, 95%) were purchased from Sigma-Aldrich (Massachusetts, USA). Hexane (Optima), chloroform (HPLC grade), ethyl alcohol (anhydrous, >95%), high purity cadmium standard (1,000 ppm, in 5% HNO_3_), high purity lead standard (10,000 ppm, in 5% HNO_3_), and nitric acid (trace metal grade) were purchased from Fisher Scientific (New Hampshire, USA). 1,2-distearoyl-sn-glycero-3-phosphoethanolamine-N-[amino(polyethylene glycol)-2000] (DSPE-PEG_2k_) was purchased from Avanti Polar Lipids (Alabama, USA).

### Synthesis of PbS cores

The synthesis of PbS QDs was adapted from previously published methods.^35,44^ For the medium (M) and large (L) sized cores, 5 g (18 mmol) of PbCl_2_ was dissolved into 14 mL OlAm by heating at 160°C for 1 hour until fully dissolved. The solution was cooled to 120°C, degassed for 30 min, and backfilled with argon. In the air-free glovebox, 76 mg (2.375 mmol) of S was dissolved in 2 mL OlAm to prepare the anion precursor. Once the PbCl_2_/OlAm solution stabilized at 100°C, the S/OlAm solution was bolus injected into the reaction and the reaction temperature maintained for 7.5 min and 30 min to generate M and L cores, respectively. Rapid quenching of the reaction via an injection of 30 mL liquid nitrogen-chilled ethanol:hexane (2:1 ratio) guaranteed the homogeneity of the particles by dropping the reaction temperature below 10°C, hindering additional growth. For the S core, the precursor concentration was reduced to 0.834 g (3 mmol) PbCl_2_ in 7.5 mL OlAm and 24 mg (0.75 mmol) S in 2.25 mL OlAm and reaction proceeded at 80°C for 7.5 minutes before rapid quenching. The reduced precursor concentration and lower temperature are used to reduce the particle nucleation and growth rates to increase homogeneity control for smaller particles.

To increase the stability of the as-synthesized particles, they were ligand exchanged to generate a more stable oleic acid corona. In an argon-filled glove box, the reaction solution was centrifuged and the pelleted particles resuspended in 50 mL hexane:OA (1:4) 50 mL with vigorous shaking. The mixture was then precipitated via centrifugation with OA acting as anti-solvent and the pellet again resuspended in hexane/OA mixture.^34^ The particles were resuspended and pelleted 3-5 times to ensure adequate ligand exchange. Initially the supernatant was dark red and turbid; the cleaning/ligand exchange process was complete when the supernatant appeared transparent with slight brown color. The final product was resuspended in 10 mL hexane, centrifuged to remove any additional insoluble lead precursor, and stored in argon filled glove box until further usage.

### Synthesis of PbS/CdS core/shell particles

Cadmium oleate (CdOA_2_) in ODE was used as cation precursor to facilitate the cation exchange for the core/shell synthesis. Specifically, 0.2 M cadmium oleate was prepared by dissolving 2.57 g (20 mmol) CdO in 28 mL (80 mmol) OA and 72 mL ODE. The solution was heated to 240°C until it clarified, cooled to 120°C, and degassed for 1 hr before backfilling with argon. The mixture was stored under argon until use. For each cation exchange reaction, 20 mL of 0.2 M CdOA_2_/ODE was heated to the reaction temperature (120°C for QD1100 and QD1300; 90°C for QD1550) before 3 mL of cleaned PbS cores in hexane was added for a 30 min anneal/reaction. After the reaction, the QD solution was diluted with 1× hexane and 3× ethanol by volume and pelleted by centrifugation. The procedure was repeated three times to remove excess ligand and precursor. The final product was resuspended in chloroform and stored in the fridge until needed.

### Optical characterization

QDs were precipitated with hexane:ethanol (1:3) and resuspended in TCE for optical characterization; TCE was chosen to minimize solvent absorbance in the SWIR.^13,45^ Absorbance spectra of the QDs were measured with a Jasco V-770 UV-Vis-NIR spectrophotometer. Photoluminescence (PL) spectra of the core/shell QDs were recorded with a FLS1000 photoluminescence spectrometer (Edinburgh Instruments, UK) equipped with a near infrared photomultiplier tube (NIR-PMT) with sensitivity up to 1700 nm. The same spectrometer was also used to measure the absolute quantum yield (QY) values of the PbS/CdS QDs. Cuvettes containing dilute QD samples (∼0.5 abs at excitation) in TCE were placed in an integrating sphere and excited with a Xenon arc lamp excitation source with 10 nm slitwidth at 808 nm, selected to match the excitation wavelength used in the imaging studies. With pure TCE in the same sample holder used in the background scan, the absolute QY was calculated with Fluoracle® (Edinburgh Instruments, UK) software.

### Structural characterization

QDs suspended in hexane were dropcast onto a zero-background silicon sample holder for X-ray diffraction (XRD) measurements. A Burker D2 Phaser analyzer with a coupled theta-2 theta scan was used to record the diffraction profiles of different particles. XRD data were analyzed with an open-source software Fityk and the reference lines of PbS (ICDD No: 01-077-0244) and CdS (ICDD No: 01-089-0440) were obtained from the International Centre for Diffraction Data (ICDD) database.

Transmission electron microscopy (TEM) images were recorded to examine the morphology and size distribution of the particles. To prepare the samples, QDs were washed with hexane/ethanol mixture multiple times before resuspension in hexane to remove excess ligands. Then, QDs in hexane were dropcast onto ultrathin carbon film-coated copper TEM grids. After the hexane evaporated, the grids were washed with drops of acetone and then ethanol before overnight storage in a dry box. A Tecnai Osiris TEM (FEI, USA) was used for the measurements. TEM and high-resolution transmission electron microscopy (HRTEM) images were recorded with 300 kV electron beam. A selected few with 620 kx magnification were presented in this study.

For small-angle X-ray scattering (SAXS) measurements, 1.5 mm diameter disposable quartz capillaries were filled with QDs diluted in hexanes and sealed using a capillary wax applied using a battery-operated hot wax pen to provide an airtight seal once cooled. Aliquots were analyzed using a Bruker N8 Horizon, with Cu Kα radiation at 50 kV and 1000 μA. Data analysis was performed on Bruker’s DIFFRAC.SAXS software assuming a spherical particle shape with a log–normal size distribution for PbS core samples, and a spherical core/shell particle structure with an input of core-to-shell density ratio of 1.58 to represent the density differences between PbS and CdS for the core/shell samples.

The hydrodynamic diameter and zeta potential of micelle encapsulated QDs were analyzed on a Litesizer DLS 500 (Anton Paar, Austria). 1 mL sample was filtered through a 0.1 μm syringe filter and loaded into a plastic cuvette. Each measurement was repeated 3 times.

Microwave plasma atomic emission spectroscopy (MP-AES; Agilent 4210) was used to determine the elemental concentration of the particle solutions and estimate the core/shell morphology. Hydrophobic particles were precipitated with hexane/ethanol and the resulting QD pellet dissolved with trace metal grade nitric acid before dilution to 5% nitric acid with ultrapure water. Aqueous QD samples were dissolved with a 10-fold excess of pure nitric acid, then diluted with ultrapure water to 5% nitric acid.

Sample concentrations were determined using a standard curve generated by high quality elemental standards of Cd and Pb. Core radius and shell diameter were calculated by using the total diameter sizing information from SAXS measurements, combined with the molecular mass and density differences between CdS and PbS, and the relative concentrations of Cd and Pb ions found in the dissolved QD samples.

### Micelle encapsulation

For water solubility and use in the biological environment, QDs were encapsulated in DSPE-PEG_2k_ micelles following a previously published protocol.^46^ Briefly, QDs were resuspended in ∼10 mL chloroform together with a 4.5-fold mass of DSPE-PEG_2k_. The mixture was evaporated with a rotary evaporator at 65°C. Ultrapure water pre-heated to the same temperature was added to the flask together with two clean glass marbles. After vigorous swirling, the solution was passed through a 0.22 μm syringe filter to remove any aggregates. The resulting particles were centrifuged at 75,000× *g* for at least 48 hours to remove encapsulated QDs (pellet) from excess lipids and empty micelles that remained in the supernatant. The pellet was resuspended in sterile pH 7.4 phosphate-buffered saline (PBS) and stored at 4°C until further use.

### In vitro QD imaging

The IR VIVO imager is equipped with three excitation laser sources at 670, 760, and 808 nm; a volume Bragg grating for hyperspectral filter cubes; multiple bandpass, longpass, and shortpass filters on two filter wheels enabling for filter combinations; and an InGaAs camera with sensitivity up to 1600 nm operating at -80°C. TCE-suspended QDs were dropcast on a background free substrate (Kimwipe tissue), and micelle encapsulated QDs were transferred into 2 mL centrifuge tubes before they were imaged.

Specifically, the images were acquired with 808 nm excitation and hyperspectral filters from 1000 nm to 1600 nm with 2 nm step size. We recorded the particle free background with the same setting and presented the background subtracted data. Following hyperspectral imaging, the QDs solutions were diluted to be brightness matched based on their peak intensity for imaging with 6 OD-4 bandpass filters centered at 1100 nm, 1150 nm, 1300 nm, 1350 nm, 1500 nm and 1550 nm with 50 nm bandwidth. Dark count-subtracted data were used for analysis.

### Live animal imaging

BALB/c female nude mice from Charles Rivers Laboratories aged 3 to 4 weeks were purchased for *in vivo* imaging studies. Due to the autofluorescence signals caused by regular diet, the mice were fed a purified OpenStandard Diet without dye (D11112201N) from Research Diets, Inc. to minimize the autofluorescence background signal.^6^ All procedures involving animals were reviewed and approved by the Institutional Animal Care and Use Committee of Boston University (IACUC Protocol No. PROTO201800466).

To examine lymphatic drainage with SWIR imaging, QD1550 was selected for time-lapse lymphatic imaging. 50 μL of 8 μM of QD1550 in PBS were subcutaneously injected at the tail base of the mouse. For this imaging study, 808 nm excitation laser at 1.6 mW/mm^2^ was used with 1250LP and 1550BP filters with 0.1 s and 0.5 s exposure, respectively. Background subtracted images were captured every 90 seconds from roughly 3-min post-injection (p.i.) to 30-min p.i., and from 90-min p.i. to 130-min p.i. Images were analyzed with ImageJ. Curving fitting of the data was completed with OriginPro 2022.

Based on the lymphatic imaging study, three-colored multiplexed imaging was demonstrated with the lymphatic pathway and vasculature imaging. QD1100 and QD1300 were used to map out lymphatic structures while QD1550 was used to image the vasculature. With brightness matched dilution samples from *ex vivo* imaging, 50 μL of 1.2 μM QD1100 was subcutaneously injected to the left side of the tail base (with mouse in supine position) and 50 μL of 0.7 μM QD1300 was similarly injected to the right side of the tail base. The lymphatic drainage was allowed for 2 hr while images was acquired with 1100 nm, 1150 nm, 1300 nm, 1350 nm bandpass filters under 1.6 mW/mm^2^ 808 nm laser excitation for 0.5 second exposure. At 2-hr p.i., 100 μL of 8 μM QD1550 was retro-orbital (RO) injected to the mouse vasculature structure. Following the injection, camera dark count-subtracted images in the same above four bandpass filters together with additional with 1500 nm, 1550 nm bandpass filters were obtained in the same setting. Mouse was also imaged with 1000 nm and 1250 nm longpass filters with 0.05 seconds exposure time at a slightly different position due to repositioning. The bandpass filter images were colored yellow, magenta, and cyan accordingly in ImageJ. The combined image with 1150 nm, 1300 nm and 1500 nm bandpass filters was composited with ImageJ in CMYK mode.

## Acknowledgement

The research reported in this publication was supported by the National Institute of General Medical Sciences of the National Institutes of Health (NIH NIGMS) (Award No. R01GM129437).

## Author Contribution

X.Z. contributed to conceptualization, experimentation, data analysis and manuscript writing. A.P. and A.M.S. contributed to material synthesis. Y.S. contributed to electron microscopy. A.M.D. contributed to conceptualization, experimentation, supervision, and manuscript writing.

