## Supplemental Info for "Multiplexed Short-wave Infrared Imaging Highlights Anatomical Structures in Mice"

**Supplemental Information**

**Figure S1. Representative TEM images**
**Figure S2. Small angle X-ray scattering (SAXS) measurements**
**Table S1. QD ensemble diameters and 1s absorbance peak positions**
**Figure S3. Dynamic light scattering (DLS) measurements**
**Figure S4. *In vitro* imaging of QD1100, QD1300, and QD1550**
**Figure S5. Additional images and analysis for time lapse lymphatic drainage imaging**
**Figure S6. Mouse lymphatic system imaged with QD1100 and QD1300 at t = 2 hour**
**Figure S7. Additional images of the three-color multiplexed imaging**

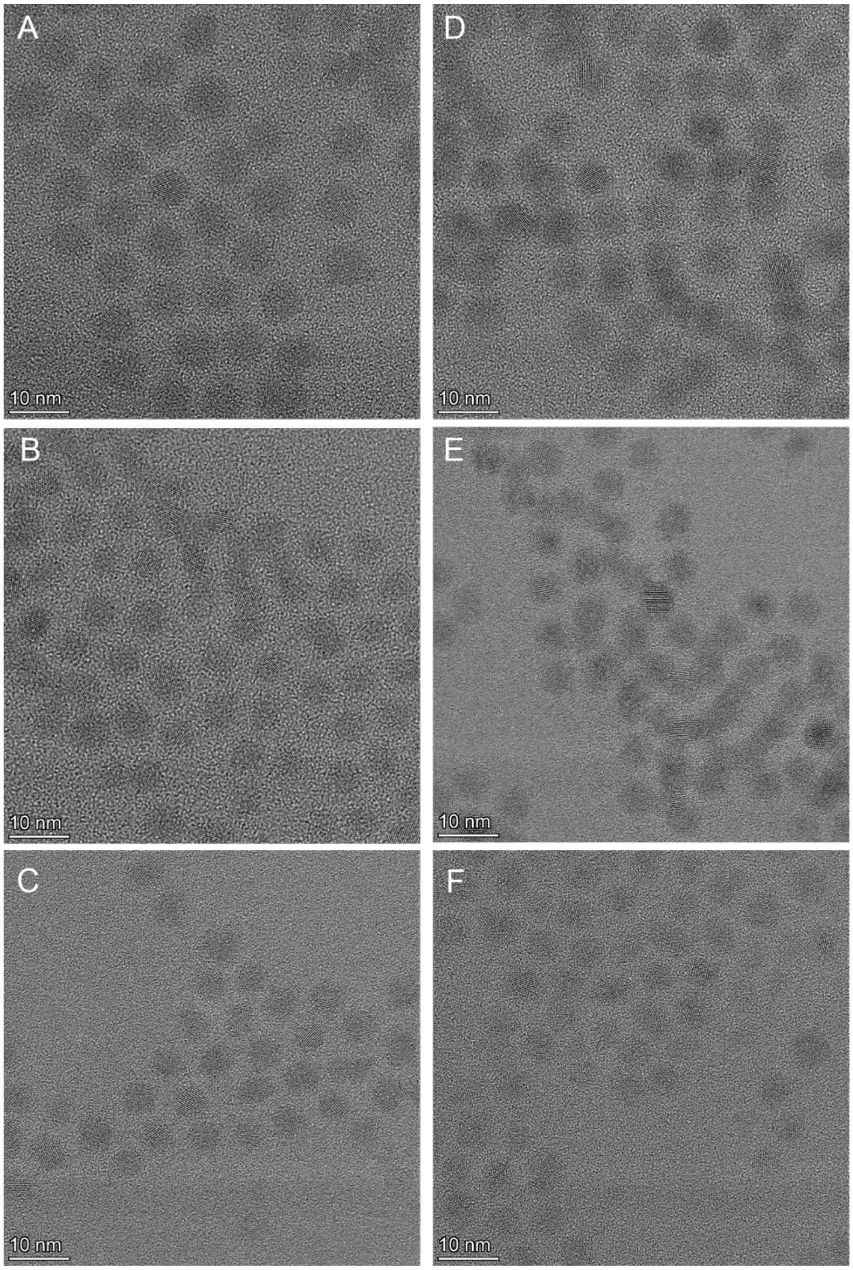


**Figure S1. Representative TEM images** of A) L cores; B) M cores; C) S cores; D) QD1550; E) QD1300; F) QD1100.

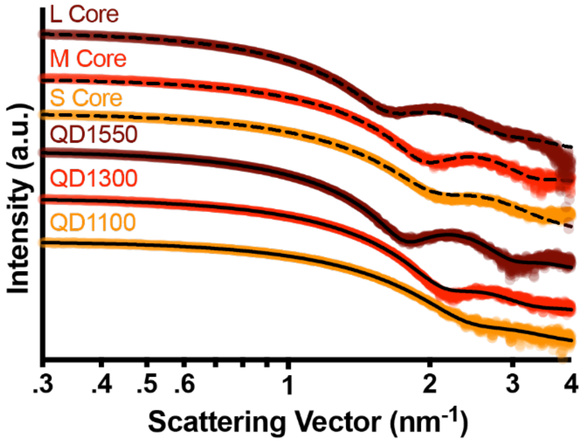


**Figure S2. Small angle X-ray scattering (SAXS) measurements** and corresponding fit curves using DIFFRAC.SAXS software (Bruker, US). Modeling assumed a spherical particle shape with a log-normal size distribution for PbS core samples and a spherical core/shell particle structure with an input of core-to-shell density ratio of 1.58 to represent the density differences between PbS and CdS for the core/shell samples.

**Table S1. QD ensemble diameters and 1s absorbance peak positions.**

| Sample | SAXS Diameter (nm) | 1s peak position (HWHM) (nm) | 1s peak position (HWHM) (eV) | PL peak position[a] (FWHM) (nm) | PL peak position[a] (FWHM) (eV) |
|---|---|---|---|---|---|
| S | 5.71 ± 0.89 | 1220 (62) | 0.98 (0.0491) | N/A | N/A |
| M | 6.29 ± 0.67 | 1370 (72) | 0.90 (0.0451) | N/A | N/A |
| L | 7.41 ± 0.98 | 1575 (66) | 0.78 (0.0371) | N/A | N/A |
| QD1100 | 5.41 ± 0.87 | 965 (84) | 1.28 (0.1026) | 1114 (192) | 1.11 (0.189) |
| QD1300 | 6.26 ± 0.82 | 1200 (78) | 1.03 (0.0632) | 1312 (182) | 0.94 (0.130) |
| QD1550 | 7.49 ± 0.70 | 1465 (72) | 0.84 (0.0397) | 1544 (186) | 0.80 (0.095) |

[a] Photoluminescence measurements taken of QDs dissolved in TCE.

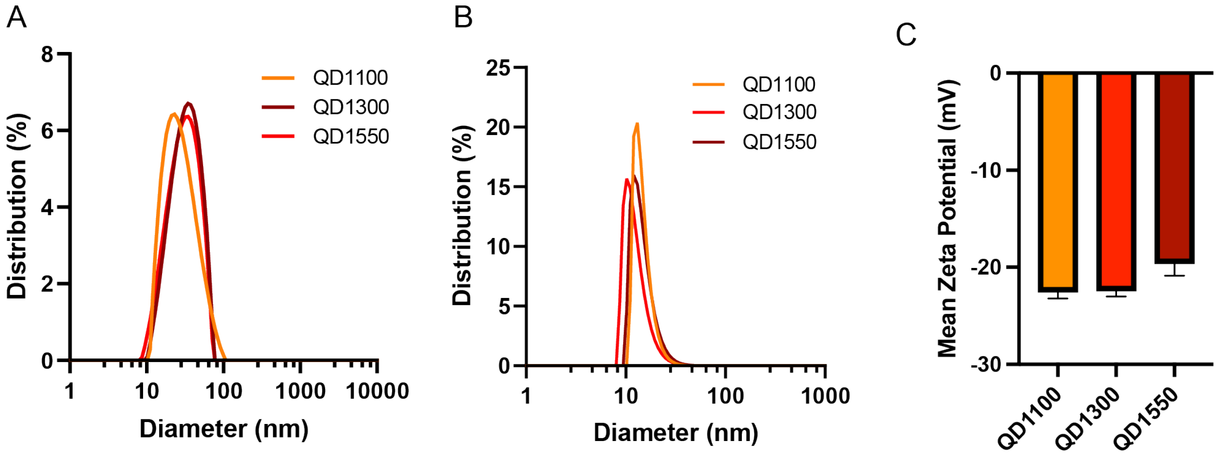


**Figure S3. Dynamic light scattering (DLS) measurements.** A) Intensity-weighted hydrodynamic diameter, B) number weighted hydrodynamic diameter, and C) Zeta potential measurements of micelle-encapsulated QDs in aqueous solution.

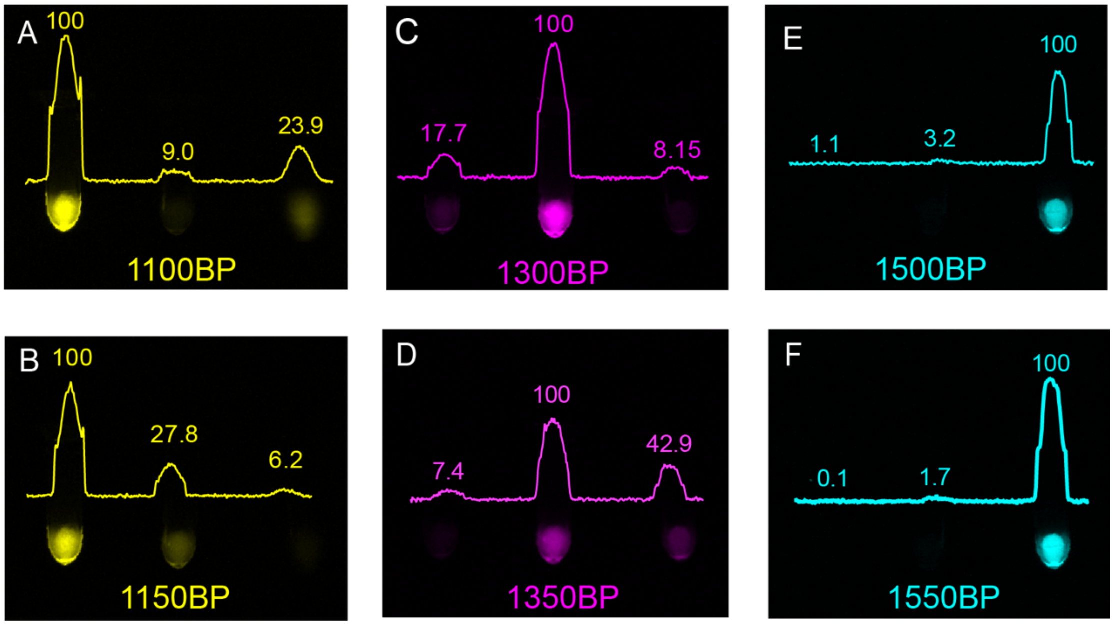


**Figure S4. *In vitro* imaging of QD1100, QD1300, and QD1550** imaged with bandpass filters. A– F) PbS/CdS QDs imaged with 1100BP, 1150BP, 1300BP, 1350BP, 1500BP and 1550BP filters. Images are pseudo-colored yellow, magenta, and cyan. Intensity linescans are normalized to the QD ensemble with maximum intensity, and the relative intensities of the other tubes are presented.

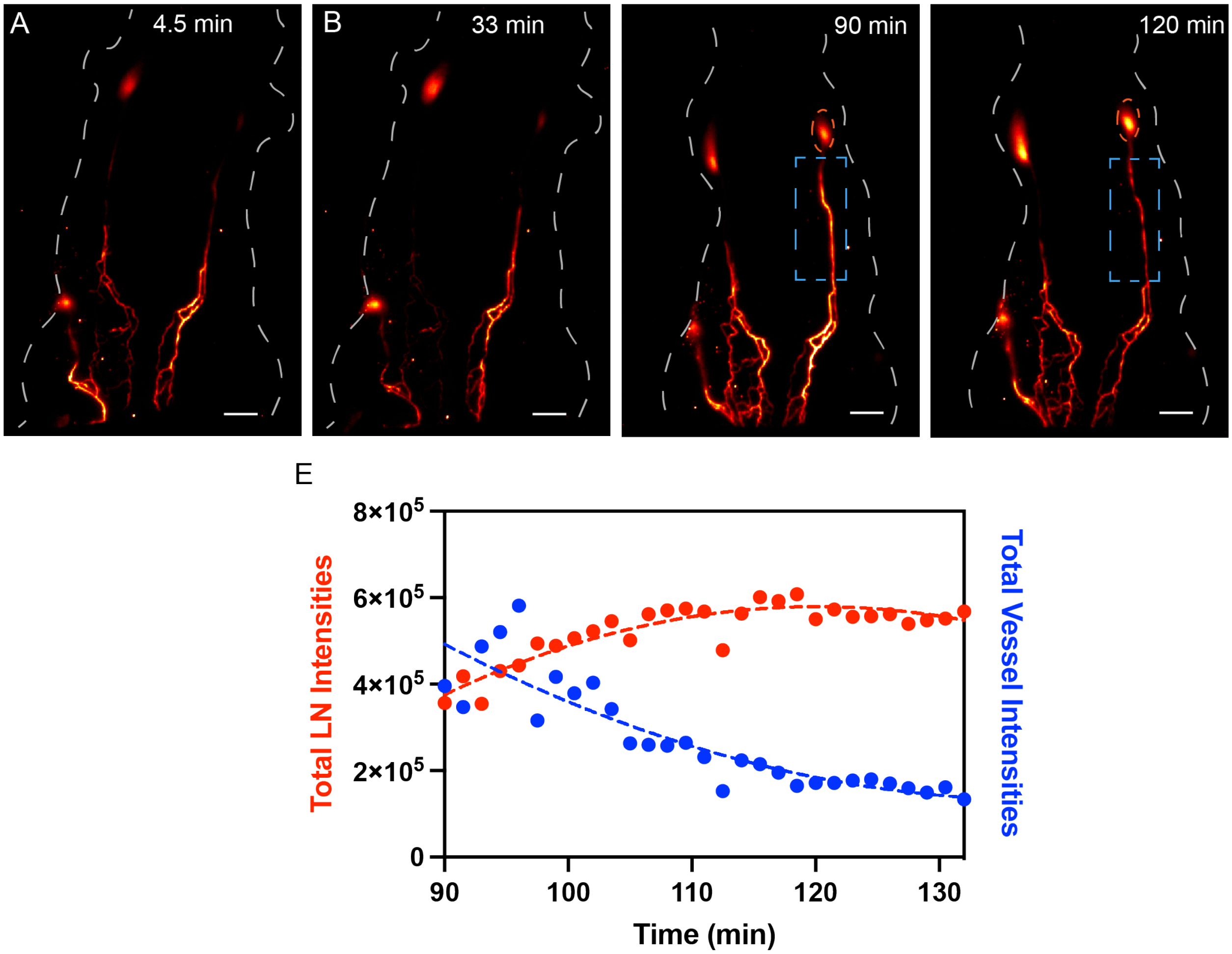


**Figure S5. Additional images and analysis for time lapse lymphatic drainage imaging.** A – D) Mouse lymphatic system was imaged with 1550BP filters at 4.5 min, 30 min, 90 min and 120 min post injection, respectively (scale bar = 5 mm). E) Total signal intensity at right axillary lymph node (oval ROI in C – D) and total signal intensity from a lymphatic vessel (rectangular ROI in C – D) from 90 minute to 133 min post injection. The data were fit with a quadratic non-linear regression model to guide the eye (dashed lines).

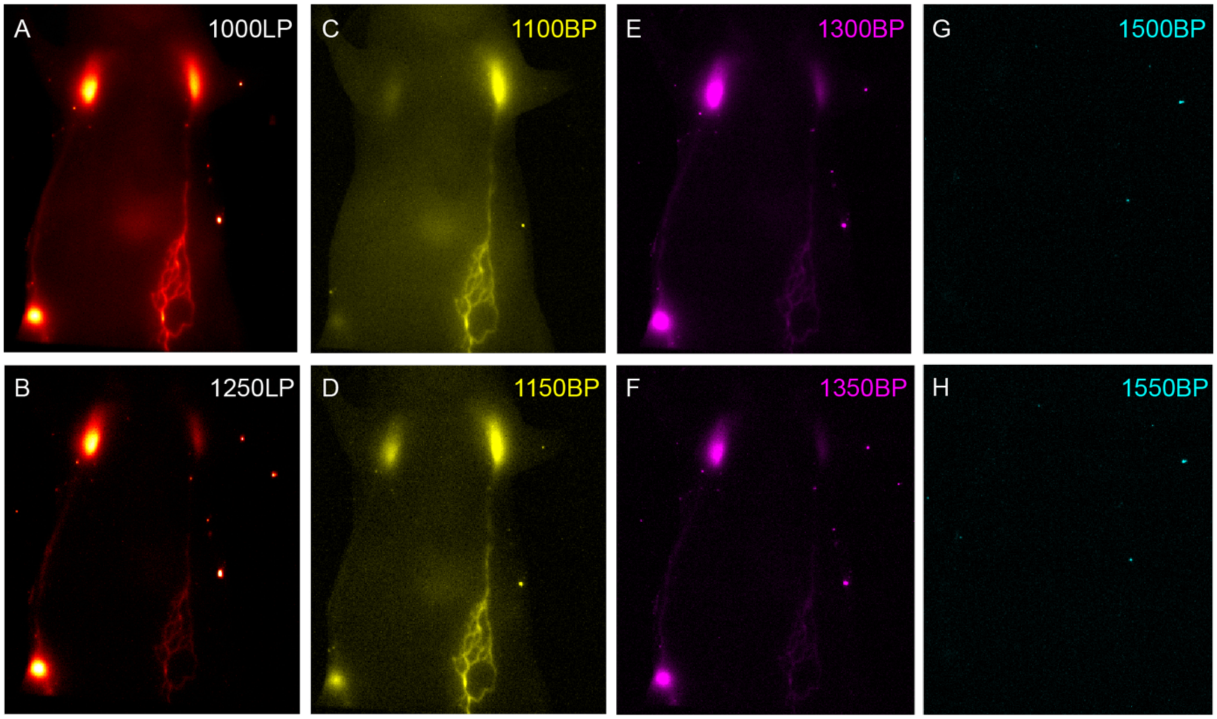


**Figure S6. Mouse lymphatic system imaged with QD1100 and QD1300 at t = 2 hour** (prior to the RO injection). A – B) Mouse imaged with 1000LP and 1250LP filters. C – H) Mouse imaged with 1100BP, 1150BP, 1300BP, 1350BP, 1500BP, and 1550BP filters.

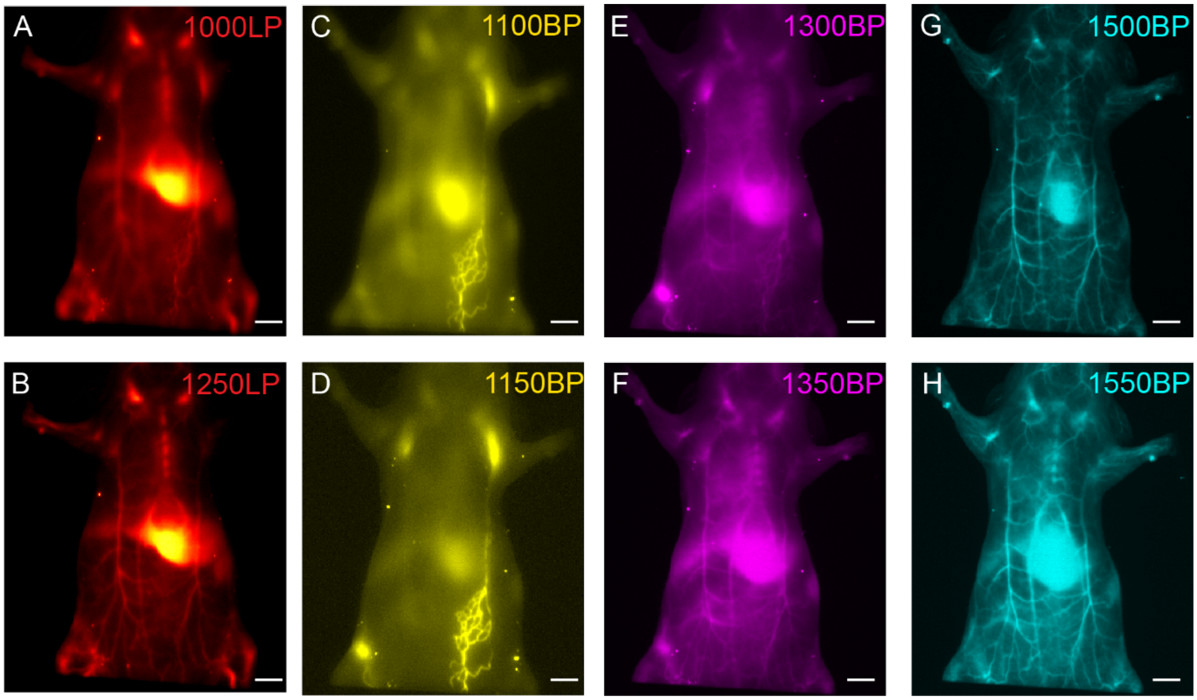


**Figure S7. Additional images of the three-color multiplexed imaging.** A – B) Mouse imaged with 1000LP and 1250LP filters. C – H) Mouse imaged with 1100BP, 1150BP, 1300BP, 1350BP, 1500BP, and 1550BP filters. Mouse positioned slightly differently in A-B compared to C-H.
